# The perception of color and material in naturalistic tasks

**DOI:** 10.1101/288662

**Authors:** David H. Brainard, Nicolas P. Cottaris, Ana Radonjić

## Abstract

Perceived object color and material help us to select and interact with objects. Because there is no simple mapping between the pattern of an object’s image on the retina and its physical reflectance, our perceptions of color and material are the result of sophisticated visual computations. A long-standing goal in vision science is to describe how these computations work, particularly as they act to stabilize perceived color and material against variation in scene factors extrinsic to object surface properties, such as the illumination. If we take seriously the notion that perceived color and material are useful because they help guide behavior in natural tasks, then we need experiments that measure and models that describe how they are used in such tasks. To this end, we have developed selection-based methods and accompanying perceptual models for studying perceived object color and material. This focused review highlights key aspects of our work. It includes a discussion of future directions and challenges, as well as an outline of a computational observer model that incorporates early, known, stages of visual processing and that clarifies how early vision shapes selection performance.

**Media Summary:** Perceived object color and material help us to select and interact with objects. There is no simple mapping between the pattern of an object’s image on the retina and its physical reflectance; our perceptions of color and material are the result of sophisticated visual computations. A long-standing goal in vision science is to describe how these computations work. We have developed selection-based methods and accompanying perceptual models for studying perceived object color and material. This focused review highlights key aspects of our work and includes a discussion of future directions and challenges.

## Introduction

Perceived object color and material properties help us to identify, select and interact with objects. Figure 1 illustrates the use of both color and material perception. To be useful, these percepts must be well correlated with the physical properties of object surfaces. Such properties are not sensed directly. Rather, vision begins with the retinal image formed from the complex pattern of light that reflects from objects to the eye. This light is shaped both by object surface reflectance and by reflectance-extrinsic factors, including the spectral and geometric properties of the illumination as well as the shape, position and pose of the objects. Because there is no simple mapping between the pattern of an object’s image on the retina and its physical reflectance properties, our percepts of object color and material are the result of sophisticated visual computations. A long-standing goal in vision science is to describe how these computations work.

**Figure 1.**
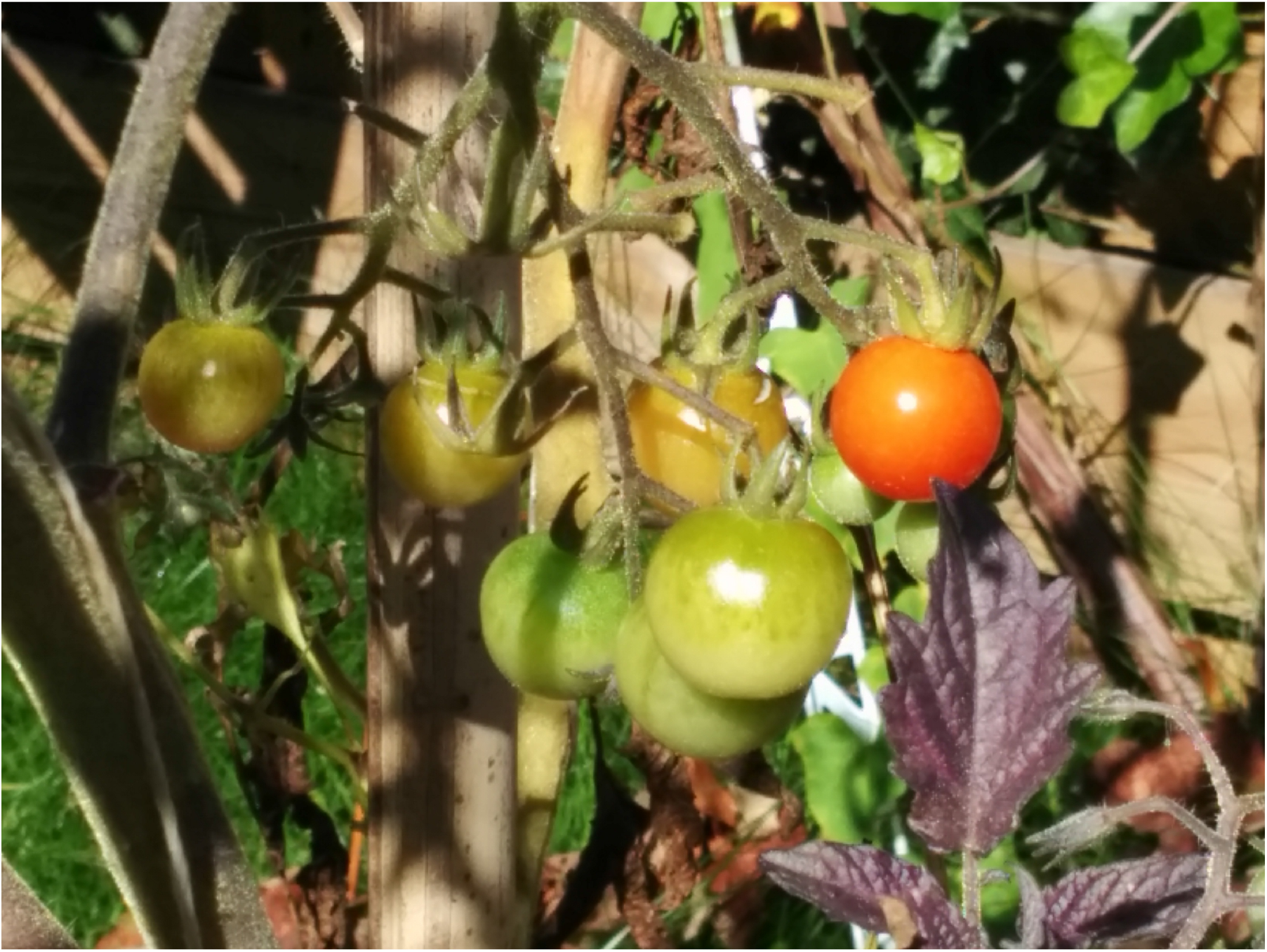
Color and material help us judge object properties. Color guides us towards the rightmost, reddest, tomato if we seek a ripe tomato for immediate consumption. If instead we are planning a dinner featuring fried tomatoes, we would select the green ones. When choosing amongst these, perceived material provides additional information, allowing us to select smoother more glossy tomatoes in preference to their rougher counterparts. Other daily life examples include identifying which of many similarly shaped cars in a large parking lot is ours and deciding where to step on an icy sidewalk. Photo courtesy of Jeroen van de Peppel.

Object color is the perceptual correlate of object spectral surface reflectance, The visual system processes the retinal image to stabilize object color against changes in the spectrum of the illumination. This is called color constancy [1-5].

Perceived material is the perceptual correlate of how object surfaces reflect light in different directions (e.g., the surface’s bidirectional reflectance distribution function, BRDF). Material perception has been the topic of energetic study over the past 15 years [6]. In parallel with color constancy, there are important questions of how vision stabilizes material percepts against variation in the geometric properties of the illumination, as well as against variation in object shape and pose.

A widely-used method for studying color constancy is asymmetric matching. Subjects adjust the color of a comparison object, seen under one illumination, so that its color matches that of a reference object, seen under a different illumination [7-9]. The correspondence in appearance established by asymmetric matching may be used to quantify the degree of human color constancy, and to study what cues are used to achieve it. Similar methods have also been used to study the stability of perceived object material [10-14].

Asymmetric matching is not a natural task. When seeking the ripest tomato (Figure 1), we do not adjust the reflectance properties of one of the tomatoes until it looks maximally tasty. Rather, we must select from a set of available choices whose properties are fixed.

If we take seriously the notion that perceived color and material are useful because they help guide selection, then we need to measure how color and material percepts guide selection, particularly in the face of variation in surface-extrinsic scene properties. We have developed such selection-based methods and corresponding perceptual models. This focused review highlights key aspects of our work. It includes a discussion of future directions and challenges, as well as an outline of a computational observer model that incorporates early, known, stages of visual processing and that clarifies how early vision shapes selection performance. Fuller descriptions of the experiments, perceptual models and main results are available in our published papers [15-18].

## Selection-Based Color Constancy

We began our study of selection-based color constancy with the stimuli and task illustrated in Figure 2. On each trial, subjects viewed a computer rendering of an illuminated cube. On the cube were three “buttons”: a reference button on the right and two comparison buttons on the left. The subject’s task was to indicate which of the comparisons was most similar in color to the reference.

The rendered scene contained two different light sources, which allowed us to manipulate the illumination impinging on two faces of the cube. Figure 2 illustrates a case where the comparison buttons’ illumination is bluer than the reference button’s. Thus, subjects had to compare color across a change in illumination.

**Figure 2.**
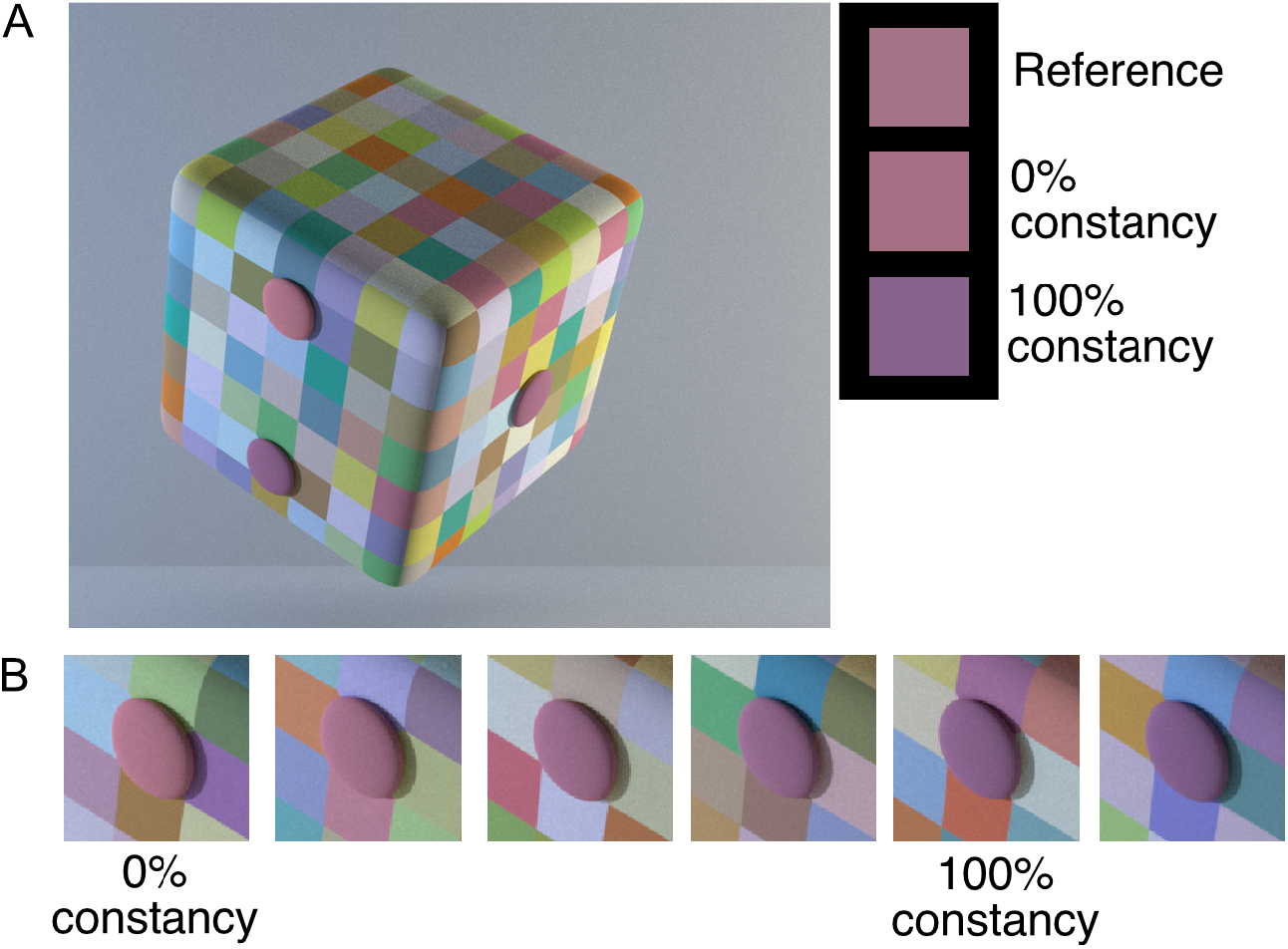
Basic color selection task. **A)** Subjects viewed a computer rendering of a cube with reference button on the right and two comparison buttons on the left. The illumination differed across the cube. The 0% (upper) and 100% constancy (lower) comparisons are shown. The squares on the right have RGB values matched to the reference, upper comparison, and lower comparison. Panel A is adopted from Figure 1 of [15] © ARVO. **B**) For a given reference, there were six possible comparisons. Panel B is reproduced from Figure 2 of [15] © ARVO.

For each reference, subjects judged all pairwise combinations of six comparisons (Figure 2B). Two of the comparisons provide conceptual anchors. One represents 0% constancy. It reflected light that produced the same excitations in the visual system’s three classes of cones as the light reflected from the reference. To produce the 0% constancy comparison, we rendered a button with a different surface reflectance than that of the reference, to compensate for the difference in illumination across the cube. The 0% comparison would be chosen as identical to the reference by a visual system that represented color directly on the basis of the light reflected from the buttons. This is illustrated in Figure 2A, where spatially uniform patches that have RGB values matched to the center of each button are shown. When seen against a common background, the patches from the reference and 0% constancy comparison match.

The 100% constancy comparison was generated by rendering a button with the same reflectance as the reference. This reflected different light to the eye than the reference, because of the change in illumination (lower square patch on the right of 2A). The 100% constancy comparison would be chosen by a visual system whose color representation was perfectly correlated with object spectral reflectance.

Three other members of the comparison set were chosen to lie between the 0% and 100% constancy comparisons. Selection of these indicates partial constancy, with the degree corresponding to how close the choice was to the 100% constancy comparison. The remaining comparison, shown at the right of 2B, represents ‘over-constancy’, that is compensation for a difference in illumination greater than that in the scene.

Data from the basic selection task for one subject, reference, and illumination change are shown in Figure 3A. The data matrix gives selection percentages for all comparison pairs. To interpret the data, we developed a perceptual model that infers perceptual representations of object color and then uses these to predict subject’s selection behavior. The model builds on maximum-likelihood difference scaling [MLDS, 19, 20] and combines key features of multi-dimensional scaling [MDS, 21] and the theory of signal detection [TSD, 22]. As with MDS, the data are used to infer the positions of stimuli in a perceptual space, with predictions based on the distances between representations in this space. As with TSD, perceptual representations are treated as noisy, so that the model accounts for trial-to-trial variation in responses to the same stimulus.

**Figure 3.**
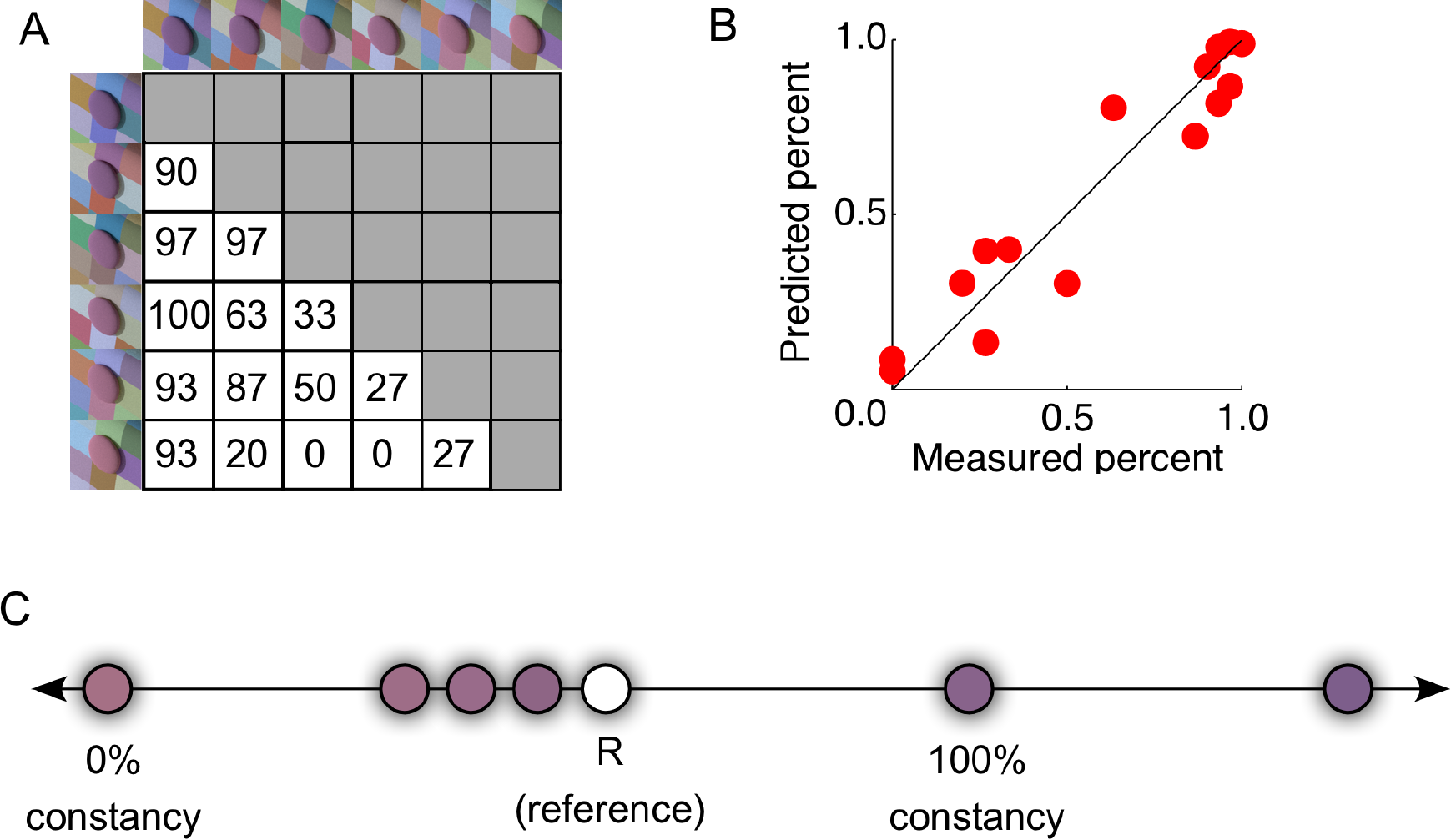
Data and model for basic color selection task. **A**). Data matrix. Rows and columns correspond to the ordered set of comparisons. Rows run from the over-constancy comparison (at top) to the 0% constancy comparison (at bottom). Columns run in the same order from left to right. Each matrix entry gives the percentage of time the comparison in the row was chosen when paired with the comparison in the column. **B**) Perceptual model concept. Each of the comparisons and the reference are positioned along a unidimensional perceptual representation of color. The positions represent the means of underlying Gaussian distributions. The positions shown represent the model fit to the data in A. **C**) Quality of the model fit to the data in A. The x-axis indicates measured selection percentage and the y-axis indicates corresponding model prediction.

Figure 3B illustrates the perceptual model. On each trial, the reference and two comparisons are assumed to be represented along one-dimension of a subjective perceptual space. Variation along this dimension represents variation in object color appearance. Because there is trial-by-trial perceptual noise, each stimulus is represented not by a single point but rather as a Gaussian distribution, with a stimulus-specific mean. The standard deviations of the Gaussians were set to one, which determines the scale of the perceptual representation.

The model predicts performance on each trial by taking a draw from the reference distribution and from the distributions for the two comparisons. It predicts the subject’s selection as the comparison whose position on that trial is closest to that of the reference. Over trials, this procedure leads to a predicted selection percentage for each member of each comparison pair.

We fit the model to the data via numerical parameter search, finding the perceptual positions that maximized the likelihood of the data. Figure 3C shows the quality of fit to the data in 3A.

The positions inferred by the model for the reference and the comparisons lie in a common perceptual space. We also have a physical stimulus description for each of the comparison stimuli. We use the corresponding stimulus and perceptual positions together with interpolation to establish a mapping between the physical and perceptual representations for the comparison stimuli. This in turn allows us to determine what comparison, had we presented it, would have the same perceptual position as the reference. We call this the selection-based match. Conceptually, the selection-based match is the comparison that would appear the same as the reference. It is also the comparison that would be chosen the majority of the time, when paired with any other possible comparison.

From the selection-based match, we compute a color constancy index (CCI) by determining where the selection-based match lies along the line between the 0% and 100% constancy comparisons, using the CIELAB uniform color space [23] to specify colorimetric coordinates. Figure 4A shows the constancy indices for the cube stimuli. The mean index was 0.47: for these stimuli the visual system adjusts for about half of the physical illumination change. The CCI varied across subjects, suggesting the possibility of individual differences in color constancy.

**Figure 4.**
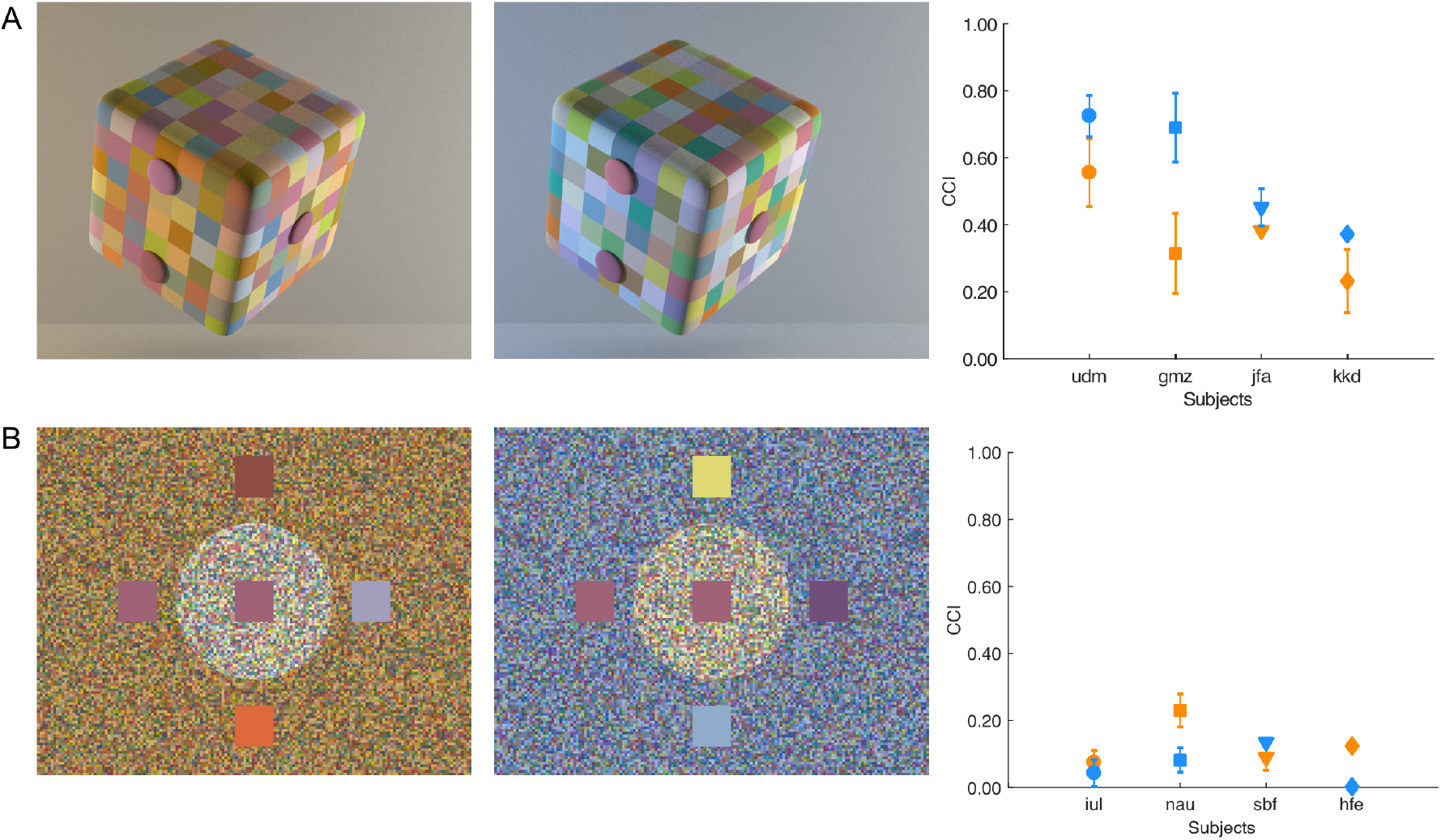
Constancy indices from the basic color selection task. **A**) The plot shows constancy indices for the naturalistic cube stimuli, for four individual subjects (x-axis). Each plotted point represents the mean CCI (y-axis) taken over four reference stimuli, with error bars showing +/-1 SEM. Blue symbols show data for a neutral to blue illuminant change (cube shown in the center). Orange symbols show data for a neutral to yellow illuminant change (cube shown at left). The initials used to track individual subject data here and elsewhere in the paper are randomly generated strings. Panel A is reproduced with permission from Figures 1 and 4B of [15] © ARVO. **B**) The plot shows constancy indices for a simplified stimulus configuration. The image at left illustrates the neutral to yellow illuminant change, while the image in the center illustrates the neutral to blue illuminant change. The reference is the central square. The two comparisons had a color appearance similar to the reference and were displayed outside the central region. Two additional distractor squares were also presented, but were rarely chosen by subjects. Panel B is reproduced with permission from Figures 5 and 6B of [15] © ARVO.

We used the basic color selection method to compare constancy obtained with the naturalistic renderings of three-dimensional cubes and for a simplified condition where flat matte stimuli were shown (Figure 4B). These simplified stimuli are similar in spirit to the “Mondrian” stimuli used in many studies of color constancy [e.g. 8, 9, 24, 25], although the spatial scale of the surface elements providing the background is smaller than has typically been employed. Across the two types of stimuli, we took care to match the colorimetric properties of the simulated illuminants, surface reflectances, references and comparisons. We found greatly reduced constancy indices for the simplified stimuli, with the mean CCI dropping from 0.47 to 0.10. The large decline in CCI for the simplified stimuli highlights the important role of image spatial structure in the neural computations that support color constancy.

## Towards More Natural Tasks

The basic color selection task is simplified relative to many real-life situations and we are interested in extending the paradigm to probe selection within progressively more naturalist tasks. Our first step [16] employs a blocks-copying task (Figure 5). The task is designed after a task introduced by Ballard, Hayhoe and colleagues for the study of short-term visual memory [26, 27]. The subject views three rendered scenes, which we refer to as the Model, the Workspace and the Source. On each trial, the subject reproduces the arrangement of four colored blocks shown in the Model by selecting from the Source.

**Figure 5.**
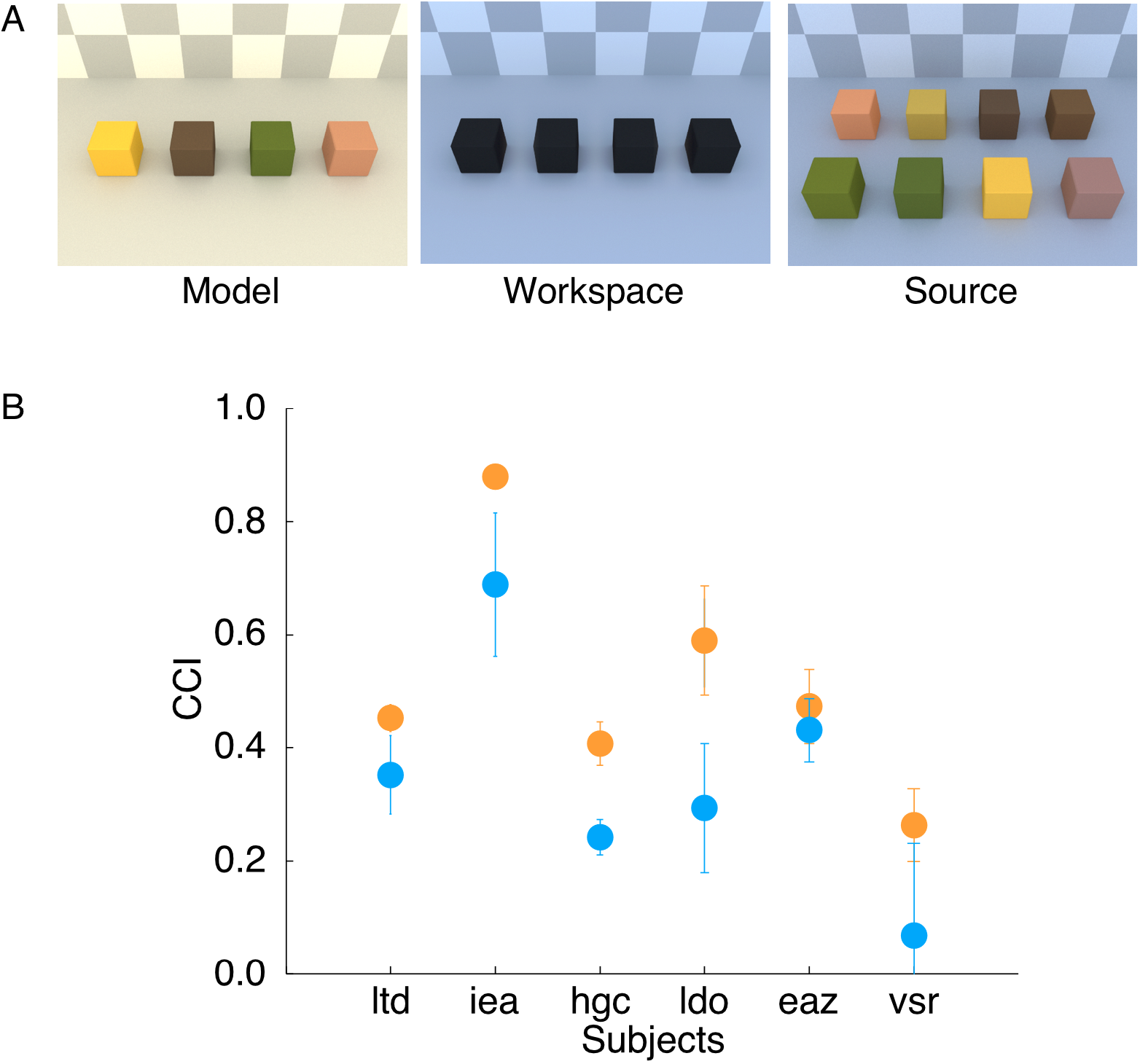
Blocks-copying task. **A**) Subjects view a computer display consisting of a Model, a Workspace, and a Source. The Model contains four colored blocks. The Source contains eight colored blocks. The subject’s task is to reproduce the Model in the Workspace. The scene illuminant varied between the Model and the Workspace/Source. Panel A is reproduced with permission from Figure 5 of [16] © ARVO. **B**) Constancy indices for the blocks-copying task. The plot is in the same format as those in Figure 4. Each point shows the mean CCI averaged over four reference blocks, with error bars showing +/- 1 SEM. Panel B is reproduced with permission from Figure 6B of [16] © ARVO.

Three features of the experimental design are worth noting. First, the four blocks in the Model are easily distinguished from each other. For each, there are just two blocks in the Source that are plausibly the same. Thus, each placement of a block in the Workspace is in essence one color selection trial. We verified that subjects rarely placed a block in the Workspace that was not one of the intended comparison pair.

Second, the Model and the Source/Workspace were rendered under different illuminants: to reproduce the Model subjects had to select blocks across a change in illumination. Across trials, subjects reproduced different arrangements of the same four blocks. Also interleaved were trials with different illumination changes.

Third subjects were instructed simply to reproduce the Model – the word “color” was not used. Rather, the use of color was invoked implicitly by the goal and structure of the task.

Because color selection is an element of the blocks-copying task, we analyzed the data using the same perceptual model we developed above. Figure 5B shows the constancy indices we obtained for six subjects, for two illuminant changes. The mean constancy index of 0.43 is quite similar to that obtained with our naturalistic cube stimuli, and again the CCI varies across subjects. This shows that the level of constancy we obtained in the basic color-selection task is maintained in a goal-directed task where the use of color is driven by task demands rather than an explicit instruction to judge similarity of color.

## Increasing the Dimensionality: Color-Material Tradeoffs

The perceptual model we developed above is based on stimulus representations in a one-dimensional space. Color vision is fundamentally trichromatic, however, and color is not the only perceptual attribute of objects. Although a one-dimensional model is a reasonable point of departure, we would like to extend the ideas to handle multiple dimensions. To that end, we studied selection where the stimuli vary not only in color but also in perceived material. We analyzed how differences in these two attributes trade off in selection.

The experimental design paralleled the basic color selection task (Figure 6A). On each trial subjects viewed a reference object in the center of the screen, flanked by two comparison objects. All of the objects had the same blobby shape and were rendered in scenes that shared the same geometry and illumination. The subject’s task was to judge which of the comparisons was most similar to the reference.

**Figure 6.**
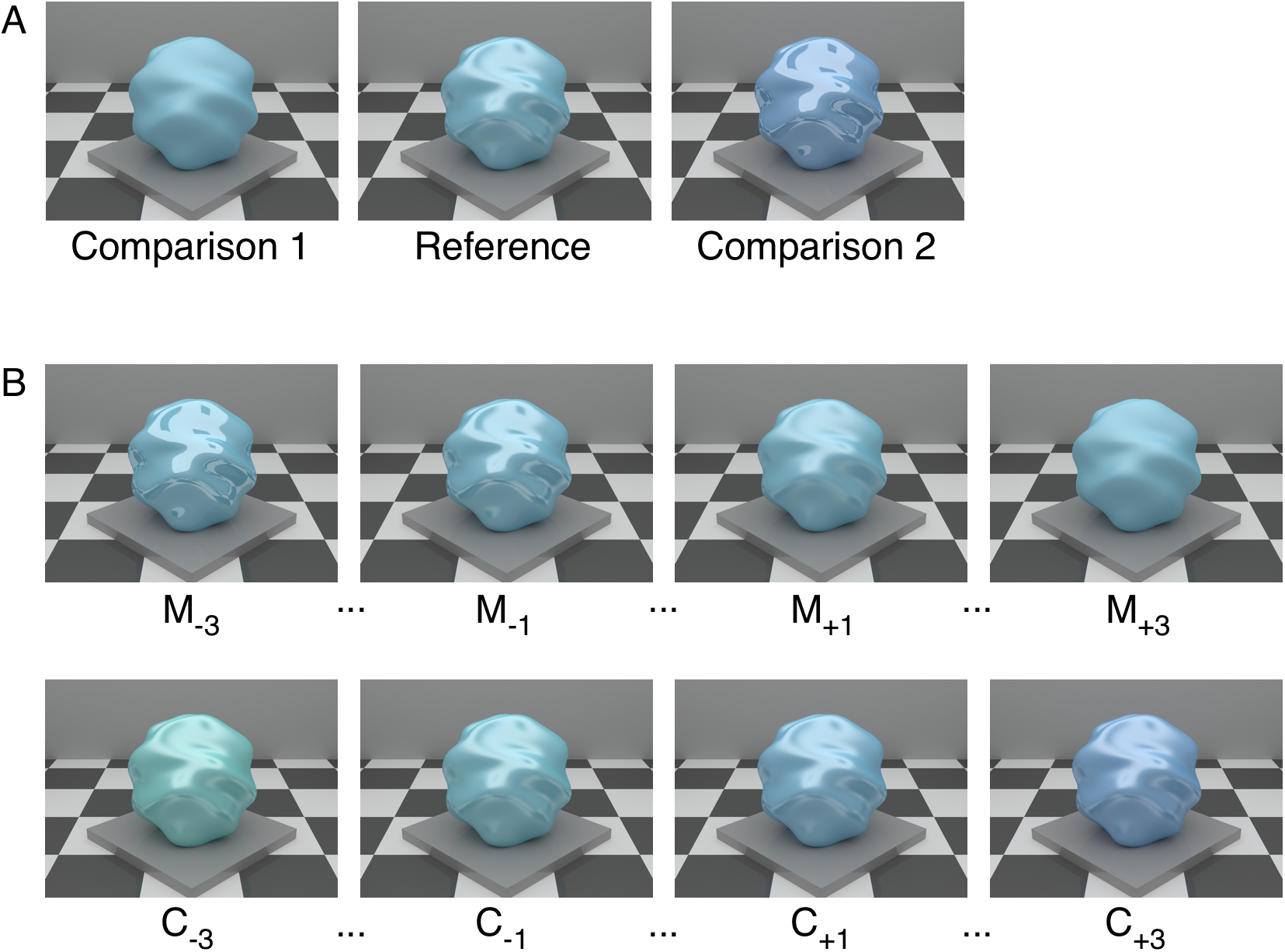
Color-material tradeoff. **A**) On each trial, the reference object was shown in the center, flanked by two comparisons. For the trial illustrated, Competitor 1 is color match M+3 and Competitor 2 is material match C+3. Panel A is reproduced from Figure 1 of [18] © IS&T. **B**) Set of comparison objects. For the 7 color matches, diffuse surface spectral reflectance is held fixed and matched to that of the reference while geometric surface reflectance (BRDF) varies. A subset of 4 color matches is shown in the top row. For the 7 material matches, surface BRDF is held fixed and matched to that of the reference, while diffuse surface spectral reflectance varies. A subset of 4 material matches is shown in the bottom row. Color match M0 was identical to material match C_0_, and both of these were identical to the reference. So, there were 13 (not 14) possible comparisons. Panel B is reproduced from Figure 2 of [18] © IS&T.

There was a single reference and 13 possible comparisons (Figure 6B), with all pairwise combinations of comparisons tested. Across the comparisons, both the diffuse spectral reflectance and geometric surface reflectance (BRDF) varied. The 13 comparison objects can be divided into two distinct sets of 7, with the reference stimulus itself being considered a member of both sets. We refer to one set as the color matches (top row of 6B). These each have the same diffuse spectral surface reflectance as the reference, but vary in their geometric surface reflectance. The color matches vary in material appearance from glossy (left of top row) to matte (right of top row).

We refer to the second set of seven comparisons as the material matches (bottom row of 6B). These have the same geometric surface reflectance function as the reference, but vary in their diffuse spectral surface reflectance. The material matches vary in color appearance from greenish (left of bottom row) to bluish (right of bottom row).

The plot in Figure 7A shows the percentage of time a color match was chosen, on trials where it was paired with one of the material matches. The x-axis gives the material match color difference. For clarity, the plot shows data for only 4 of the 7 color matches. Also not shown are data for trials where one color match was paired with another color match and for trials where one material match was presented with another material match.

**Figure 7.**
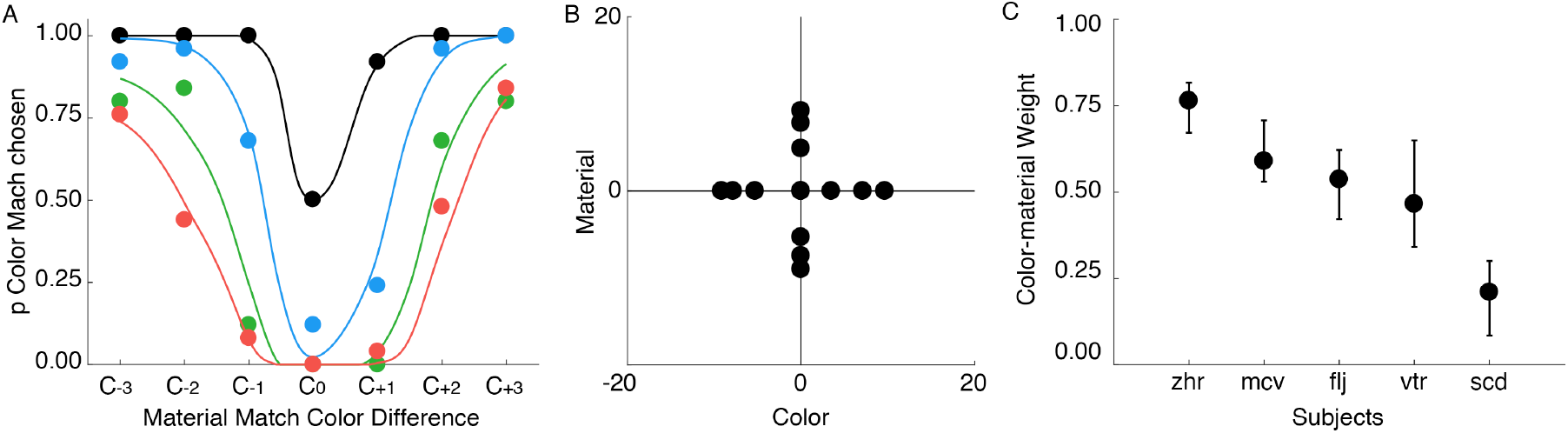
Color-material tradeoff data and model. **A**) Subset of data for one subject. For four levels of color match material difference (black, M+0; blue, M_+1_; green, M_+2_; red, M+3), the fraction of trials on which the color match was chosen is plotted (y-axis) against the material match color difference (x-axis). The black point corresponding to C_0_ and M_0_ was not measured and is plotted at its theoretical position. The smooth curves show the fit of the twodimensional perceptual model. **B**) Two-dimensional perceptual model solution for the subject whose data is shown in A. The plot shows the inferred mean positions of the stimuli in a two-dimensional perceptual space, where the x-axis represents color and the y-axis represents material. The fit is to the full data set for this subject, not just to the subset shown in A. **C**) Color-material weights inferred for individual subjects. Error bars were obtained from bootstrapping and show +/-1 standard deviation of the bootstrapped weights, plotted around the mean of the bootstrapped weights. Panel C is adopted from Figure 5 of [18] © IS&T.

Consider the black points in Figure 7A. These show trials where the color match had no material difference (M0) and was thus identical to the reference. When the material match (C_0_) was also identical to the reference, all three stimuli were identical and the point is plotted at 50%. As the color difference of the material match increases in either the positive or negative direction, the subject increasingly selects the color match. This makes sense: for these cases the color match is identical to the reference while the material match becomes increasingly distinguishable.

Now consider the red points in Figure 7A. These represent a case where the color match (M_+3_) was considerably less glossy than the reference. When the material match (C_0_) was identical to the reference, the subject always chose the material match. As a color difference in the material match was introduced, however, the subject had to judge similarity across comparisons where one differed in material and the other differed in color. As the color difference of the material match increased in either direction, the subject increasingly chose the color match. The rate of this increase is determined in part by how perceived color and material trade off in this subject’s similarity judgments.

The blue and green points show intermediate cases of color match material differences (M_+1_, M_+2_). As the material difference of the color match decreased (red -> green -> blue -> black), it took a smaller color difference in the material match to cause the subject’s selections to transition from the material to the color match. A similar pattern was seen in the data for color matches that were more glossy than the reference (M-1, M-2 and M-3, data not shown).

The data shown in Figure 7A reveal a gradual transition from material match to color match selections as a function of the material match color difference. The rate of this transition is related to how the subject weights differences in perceived color and material. The exact pattern of the data, however, also depends on the size of the perceptual differences between the stimuli. There is no easy way to equate perceptual differences between supra-threshold stimuli, and no easy way to equate the perceptual magnitude of color steps and material steps. We chose the stimuli by hand on the basis of pilot data and our own observations. The physical stimulus differences must thus be regarded as nominal, and we cannot assume that the perceptual step between (e.g.) M0 and M_+1_ is the same as that between M_+1_ and M_+2_, or between C_0_ and C_+1_. Interpreting the color-material selection data requires a perceptual model.

We developed a model that assumes that each stimulus has a mean position on two perceptual dimensions, one representing color and one representing material. To reduce the number of free parameters, we restricted the mean of the color matches to lie along the perceptual material dimension and mean of the material matches to lie along the perceptual color dimension. There was trial-by-trial variation in the perceptual position of each stimulus, and the perceptual noise was represented by two-dimensional Gaussian distributions.

Trial-by-trial predictions of the model were based on which comparison had a representation closest to the reference on each trial. We computed distance using the Euclidean distance metric. Importantly, however, we weighted differences along the perceptual color dimension by a color-material weight *w* prior to computation of distance, and differences along the perceptual material dimension by (1-w). The color-material weight served to specify how perceptual differences in color and material were combined.

We fit the model using numerical parameter search and a maximum likelihood criterion. The solution for the data set illustrated by Figure 7A is shown in 7B, with the smooth curves in 7A showing the model’s predictions. The overall spread of the stimulus representation for the two perceptual dimensions is roughly equal for this subject, but the spacing is not uniform within either dimension. The color-material weight was 0.54, indicating that this subject treated the two perceptual dimensions equally. The weight is inferred together with the perceptual positions and is an independent parameter of the model. See [18] for a fuller presentation.

We measured the color-material weight for 5 subjects. These vary considerably. Of interest going forward is to understand how stable the color-material weight for a given subject is, across variations in stimulus conditions where the relative reliability of color and material are varied. For example, if the comparisons are presented under a different spectrum of illumination than the reference, the visual system might treat color as less reliable and reduce the color-material weight.

## Discussion

### Summary

Understanding mid-level visual percepts will benefit from understanding how these percepts are formed for naturalistic stimuli and used in naturalistic tasks.

The basic color selection task [15], blocks-copying task [16], and color-material tradeoff task [18] illustrate how to use naturalistic stimuli in conjunction with naturalistic tasks to infer selection-based matches and draw inferences about perceptual abilities such as color constancy. The interested reader is directed to the original papers for more detail, and to related work by others that uses selection to investigate constancy [28-31].

### Tools to Facilitate the Use of Computer Graphics in Psychophysics

Computer graphics is important for and widely-used in visual psychophysics because it enables manipulation of the distal scene properties (e.g., object surface reflectance) with which vision is fundamentally concerned. To conduct our studies, we developed open-source software tools to facilitate the use of high-quality graphics for psychophysics. First, RenderToolbox [rendertoolbox.org, 32] allows specification of full spectral functions (in contrast to RGB values) for both object surface reflectance and illuminant spectral power distributions. In addition, it provides a convenient interface that allows Matlab access to physics-based renderers. Our recent collaborative work with the Hurlbert lab on illumination discrimination [33, 34] also takes advantage of RenderToolbox.

Second, for the blocks-copying task, we needed to re-render the spectral surface reflectance of individual blocks rapidly in response to subject selections. To accomplish this, we developed software tools for synthesizing images as linear combinations of a set of pre-rendered basis images [16, see also 35]. These tools are available in the BrainardLabToolbox (github.com/BrainardLab/BrainardLabToolbox.git).

### Perceptual Models

Interpreting selection data in a form that allows inferences about color constancy and color-material tradeoff required the development of new perceptual models. These leverage selection data to infer the perceptual representations of visual stimuli and how information is combined across perceptual dimensions. The models extend the applicability of maximum-likelihood difference scaling. The key extensions are i) for color selection, inferring the position of the reference and the comparisons within a single perceptual space and ii) for color-material tradeoff, generalizing the MLDS principles to two dimensions. Our Matlab implementation of the perceptual models is included in the BrainardLabToolbox.

There is no conceptual obstacle to extending the models to more than two dimensions. In practice, however, the amount of data required to identify model parameters grows rapidly with the number of parameters. This motivated some simplifying assumptions in our work (e.g., studying just one dimension of stimulus variation in our color selection work; assuming that the color matches have same mean perceptual color coordinate as the reference and that the material matches have the same mean perceptual material coordinate as the reference in our color-material selection work.) We view these assumptions as sufficiently realistic to allow progress, but they are unlikely to be completely accurate [36, 37]. An important goal for model dimensionality extension is to be able place stimuli within a full three-dimensional color space and infer corresponding perceptual positions. This would enable the use of selection-based methods more generally in color science.

There is need for additional work on the perceptual models. First, we would like to better characterize how effectively selection data determine model parameters. Related to this is the desire to increase the efficiency with which the measurements constrain these parameters. One approach is to develop accurate parametric models that describe the mapping between physical and perceptual representations, thus reducing the number of parameters that need to be determined. A second approach is to develop adaptive psychophysical procedures that optimize trial-by-trial stimulus selection. We have recently implemented the Quest+ adaptive psychophysical method [38] to our color-material model, and are exploring its efficacy [see also 39]. We have made freely available a Matlab implementation of Quest+ (github.com/BrainardLab/mQUESTPlus.git).

Although we have emphasized the role of the perceptual models for inferring interpretable quantities of immediate interest (the CCI; the color-material weight), these models also provide rich information about the mapping between physical and perceptual stimulus representations (Figures 3 and 7) as well about the precision of these representations [39, 40]. We think it will be of future interest to leverage the models to understand more about these aspects of performance.

### Interesting Extensions

We describe two natural experimental extensions of our work (Figures 8 and 9). The first is a variant of the blocks-copying task that could be deployed to study the role of learning via natural feedback. In a feedback variation of our experiment, the Model and Workspace are shown under a common illuminant, while the Source is shown under a different illuminant (Figure 8A). The subject moves a block from the Source to the Workspace and compares it with the Model under a common illumination, and then chooses a different block if the alternative is preferred. After experience with this version of the task, the subject is tested in the original version of the blocks task (Figure 8B) to determine the generality of learning that occurred in the feedback condition.

**Figure 8.**
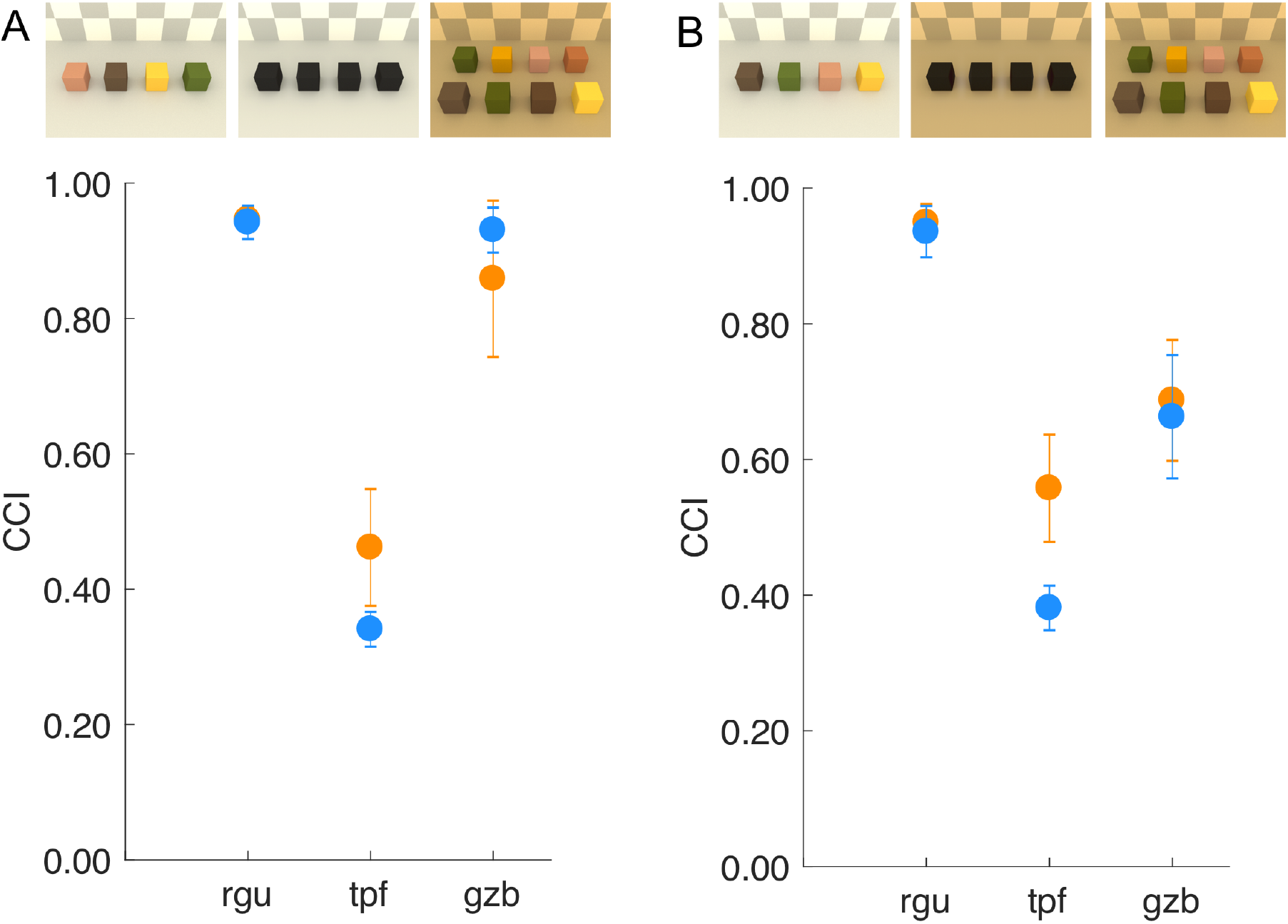
Effect of learning on constancy. **A**) Version of blocks-copying task that provides naturalistic feedback. Preliminary data from three subjects shows high CCI’s for two of three subjects in the feedback condition. **B**) When subsequently tested in a version of the task without feedback, CCI’s for one subject remained high. Data in both A and B are for two illuminant changes. For subject rgu, the data for the two illuminant changes overlap.

We collected preliminary data using this paradigm. Two of three subjects took advantage of the feedback and achieved high levels of constancy when it was available (8A). When subsequently tested without feedback, one of these subjects maintained high constancy (8B). It would be interesting to more carefully characterize these learning effects. For example, does improved constancy generalize to reference block colors and/or illuminant changes that were not presented in the feedback condition?

In many real-life scenarios involving color selection, stimuli are compared to a memory-based rather than to a physical reference. For example, when picking the most desirable tomato (Figure 1), the remembrance of tomatoes past surely plays an important role [e.g. 41, 42-46]. We have not yet introduced a memory-reference component into our selection task, but this could be done. Figure 9 illustrates one approach. The subject plays a simple adventure game. On each trial, the subject enters a room in which there are two dragons, one each from two species. The subject selects which dragon to engage. One species is friendly and the other evil, with the species distinguished by the reflectance properties of their scales. Friendly dragons give gold coins and evil dragons steal them. The subject’s task is to amass as much gold as possible. Each trial in this experiment involves a color selection and can be instrumented and analyzed using the methods described above. The key extension is that subjects must learn over trials the reflectance properties of the dragons. Early trials would be under a single illuminant. Changes in illumination would be introduced after subjects have formed a memory reference. This type of experiment could provide a fuller picture of how object color representations form and how these are combined with perception to achieve task-specific goals.

**Figure 9.**
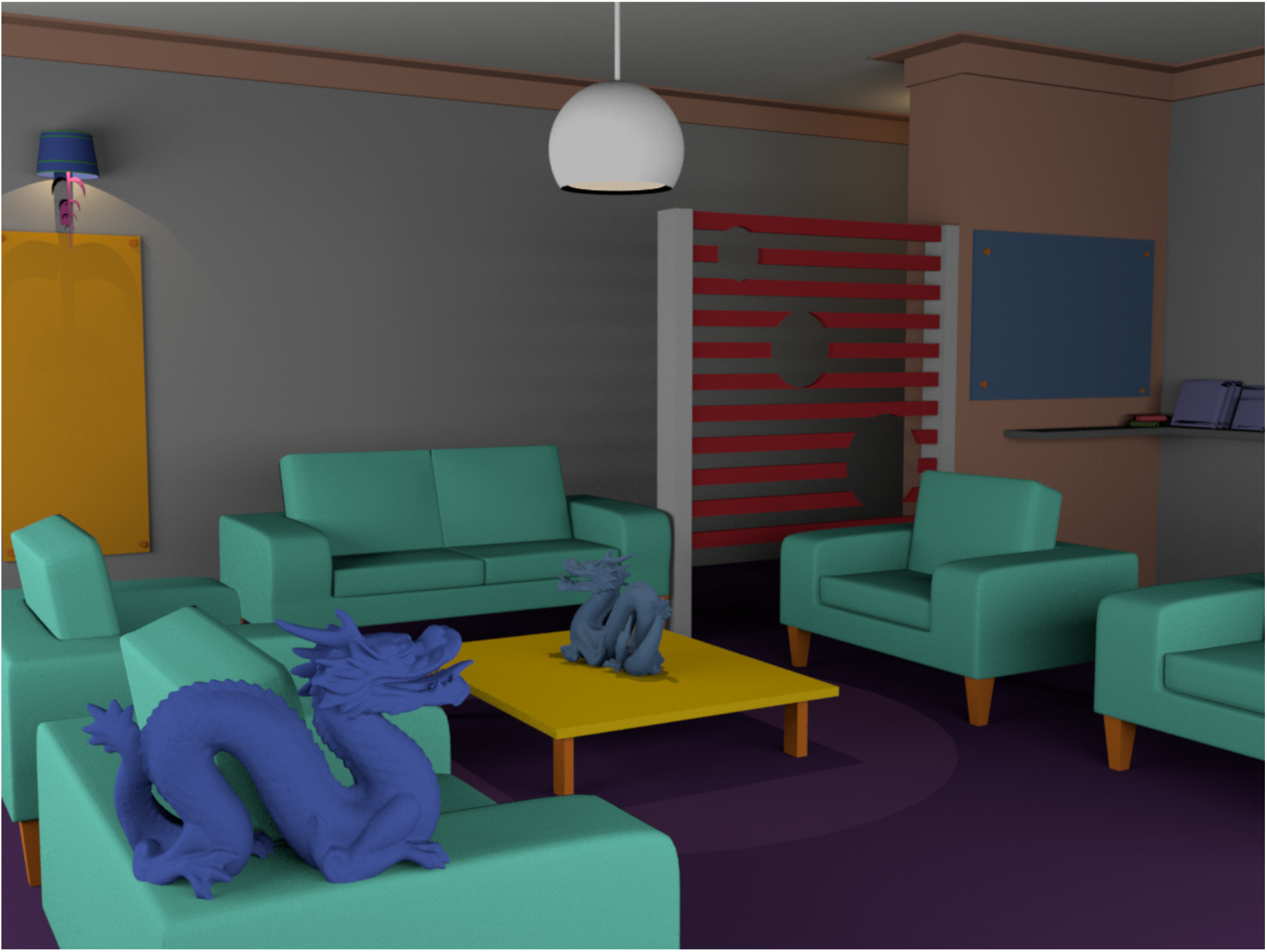
Introducing memory into the color-selection paradigm. The image depicts a possible stimulus for a color-selection adventure game. The geometry of the scene shown here (sans dragon) was created by Nextwave Media (http://www.nextwavemultimedia.com/html/3dblendermodel) and made freely available on the internet. The dragons were added and the scene rendered using RenderToolbox [32]. The appearance of the image was optimized for publication using Photoshop.

### Instructions

How subjects are instructed to judge color and lightness can affect the degree of constancy they exhibit [8, 17, 47-53]. In an asymmetric matching experiment, for example, subjects instructed to adjust a comparison so that it appeared to be “cut from the same piece of paper” as the reference showed higher constancy than those instructed more simply to match “hue and saturation” [8]. How such instructional effects should best be understood, as well as when they do and do not occur, has not been resolved [see 17 for discussion]. As we noted above, the use of color in the blocks-copying task is defined implicitly by the goal of reproducing the blocks, not by explicit reference to color. That subjects base their judgments on color follows naturally, since for these stimuli color is the only feature that differentiates individual blocks. Thus, it seems reasonable to interpret the measured CCI’s as representative of what subjects would do if performing such a task in a real setting. By asking how the brain uses color to help achieve task goals, we shift the emphasis from perception to performance and finesse the need to make distinctions between perception and cognition. The perceptual positions inferred via the perceptual model also provide the basis for task- and instructions-specific comparisons with more traditional assessments of appearance [17].

### Individual Differences

Our measurements reveal individual differences, both in the color constancy indices (Figures 4, 5, 8) and in the color-material weights (Figure 7). We do not have any direct evidence about what drives these differences. As a general matter, individual variations in performance in visual tasks have been documented and systematically studied [for a recent overview see 54]. In color perception, efforts to understand individual differences have focused on early visual processing. This work includes studies of differences in cone spectral sensitivities [55-57], relative numbers of cones of different classes [58, 59], and loci of unique hues [60]. More recently, the phenomenon of #theDress revealed striking individual differences in the perceived colors of an image of a dress, with speculation that these are related to individual differences in color constancy [61-69]. Studies that systematically investigate which individual difference factors are correlated with the constancy index differences we measure may help advance our understanding of the mechanisms of color constancy. A similar approach could be used to investigate individual differences in material perception, although these are currently less well documented.

### A Mechanistic Model

Both object color and perceived material are often described as mid-level visual phenomena, and it is thought that these judgments depend on cortical mechanisms. Even so, the signals that reach the cortex are shaped by the optics of the eye and retinal processing. These are increasingly well-understood and can now be modeled quantitatively. There is important progress to be made by understanding the role of early factors in limiting and shaping the information for performance of visual tasks, a view with roots in seminal work on ideal observer theory [70, 71]. We are developing a set of open-source Matlab tools for modeling early vision, which we call the Image Systems Engineering Toolbox for Biology [ISETBio, isetbio.org, 72, 73]. Here we illustrate how these tools may be used to understand how color-material selection would behave, were it based on the representation available at the first stage of vision.

Our stimuli were presented on a well-characterized computer-controlled monitor, so we know the spectrum at each pixel of the displayed image. Using this information, we can compute the retinal image using a model of the polychromatic point spread function (PSF) of the eye (Figure 10A). The PSF captures both optical blur due to both monochromatic ocular aberrations and to axial chromatic aberration. The retinal image is sampled by an interleaved cone mosaic consisting of L, M and S cones (Figure 10B). Each cone class is differentially sensitive to light of different wavelengths (Figure 10C).

**Figure 10.**
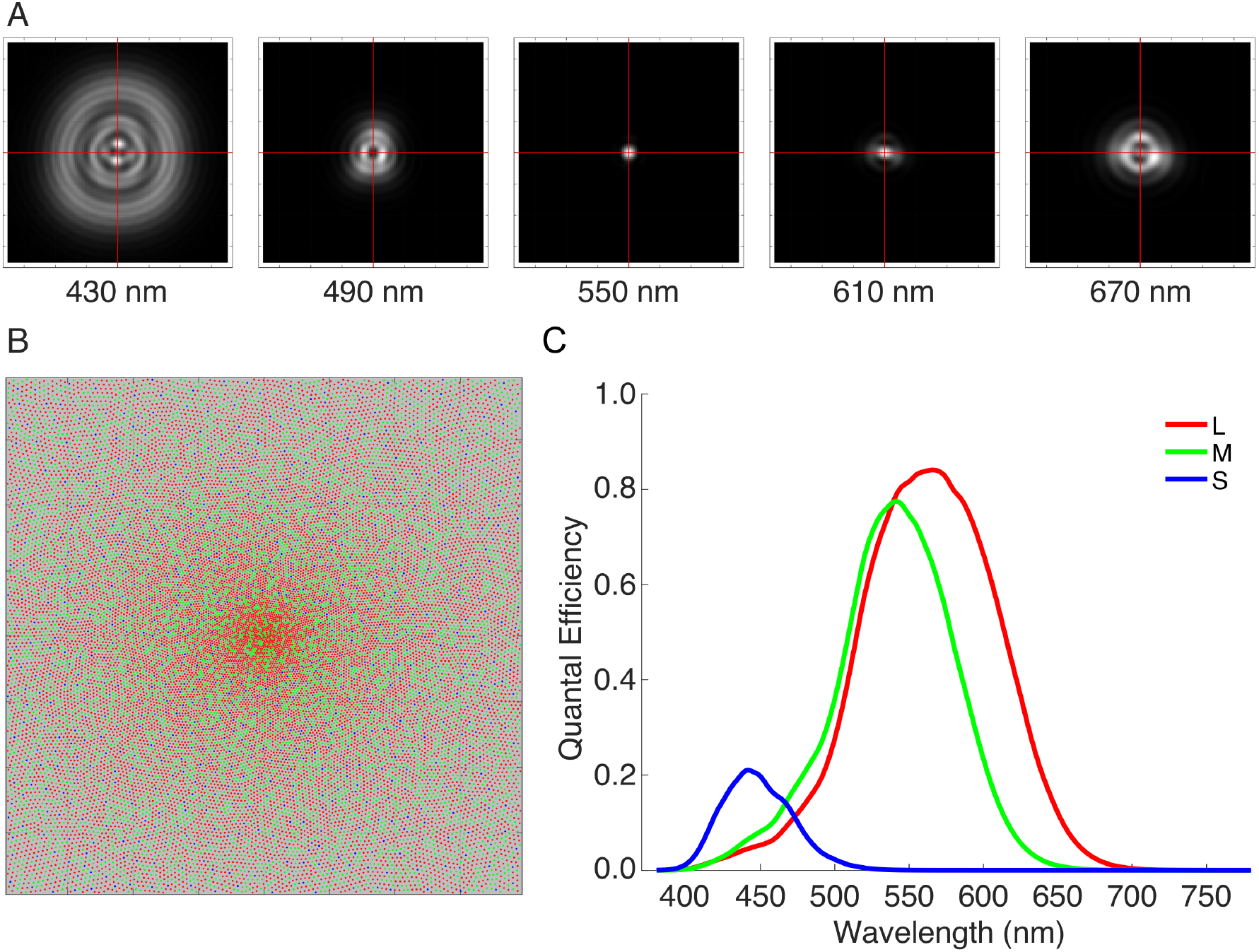
Components of ISETBio early vision model. **A**) Each image depicts an estimate of the point spread function (PSF) of the human eye for one wavelength, based on the wavefront measurements reported in [74] and assuming that the eye is accommodated to 550 nm. The effects of chromatic aberration are apparent in the broadening of the PSF away from 550 nm. Each image represents 17 by 17 arcmin of visual angle. The PSFs for 5 wavelengths are shown; calculations were done for wavelengths between 380 and 780 nm at a 10 nm sampling interval. **B**) The central portion (2 by 2 degrees of visual angle) of the interleaved retinal cone mosaic (L cones, red; M cones, green; S cones, blue). Cone density falls off with eccentricity according to the measurements of Curcio [75], and the mosaic incorporates a small S cone free central tritanopic area [76]. **C**) The plot shows the foveal quantal efficiencies (probability of isomerization per incident light quantum) of the L, M and S cones. These follow the CIE cone fundamentals [23] and incorporate estimates of cone acceptance aperture, photopigment optical density and photopigment quantal efficiency.

We computed the mean number of isomerizations to each of the stimuli we used in the color-material selection experiment for each cone in a 15 by 15 deg patch of cone mosaic, assuming a 1-second stimulus exposure. These allowed us to simulate performance in the experiment, for an observer who made selections based on the trial-by-trial similarity between each of the comparisons and the reference. Similarity was computed in the high-dimensional space where each dimension represents the isomerizations of a single cone, with smaller distance indicating greater similarity. To make predicted performance stochastic, we added independent trial-by-trial Gaussian noise to the mean number of isomerizations for each cone. The noise level was chosen by hand to bring overall performance roughly into the range of that exhibited by human subjects. Similarity was computed using Euclidean distance, and no differential weighting of individual dimensions was applied. A subset of the data obtained via the simulation is shown in Figure 11A.

**Figure 11.**
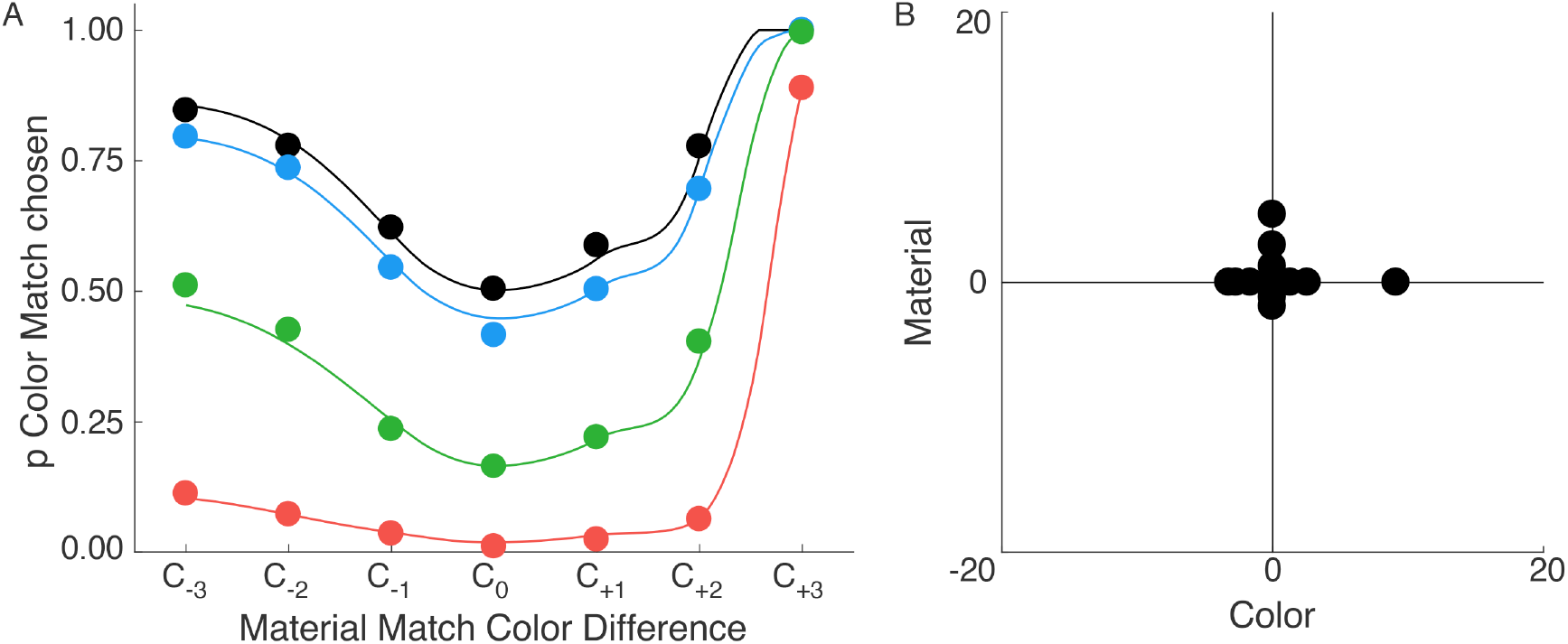
Perceptual model fit to computational observer simulations. **A**) Subset of the data obtained using the computational observer simulations. Same format as Figure 7A. **B**) Twodimensional perceptual representation obtained by fitting the data from the computational observer simulations. Same format as Figure 7B. The smooth curves in A correspond to this solution.

We fit the perceptual model to the simulated data, with the model solution shown in Figure 11B. The inferred color-material weight is 0.44. A weight close to 0.5 might be expected, since no asymmetry between color and material perception was built into the calculation. The perceptual positions, however, are distributed asymmetrically along the color and material dimensions. This differs qualitatively from the pattern we see for most subjects, where the distribution tends to be symmetric.

It is not surprising that the perceptual representation for the computational observer differs from that of human subjects: as we noted in the introduction the color and perceived material of objects are not directly available from the representation at the photoreceptor mosaic. Rather these are the result of visual computations that likely begin in the retina and continue in the cortex. For example, object material percepts are coupled with perceptual representations of three-dimensional object shape [77, 78]. That said, we do not currently know why the representation obtained for the computational observer differs from the human representation in the particular way that it does.

We plan to extend this modeling to incorporate additional known early processes, including fixational eye movements, the adaptive transformation between cone isomerization rate and photocurrent, and coding by multiple mosaics of retinal ganglion cells. To provide a good description of the early representation of color, the latter will need to include a model of the opponent combination of signals from the three classes of cones. This type of modeling has the potential to clarify which aspects of mid-level vision are follow-on consequences of early visual processing, and which require further explanation in terms of later, presumably cortical, mechanisms. One interesting question that we may be able to address is the degree to which known adaptive transformations that occur in the retina account for human color constancy.

## Acknowledgments

Supported by NIH RO1 EY10016, NIH Core Grant P30 EY001583, and Simons Foundation Collaboration on the Global Brain Grant 324759. We thank Brian Wandell for many discussions about computational observer modeling and for his lead role on the ISETBio project, as well as for providing comments on the manuscript.

## Author Contributions

The first draft of the manuscript was written by DHB. AR and DHB edited the manuscript. The bulk of the experimental work and perceptual model development is described in more detail in published papers on which AR, NCP and DHB are authors [15-18]. For these, AR and DHB conceived the experiments and perceptual models, AR and NCP implemented the experiments, AR and DHB implemented the perceptual models, and AR collected and analyzed the data. The computational observer model presented here was developed by NCP and DHB. It takes advantage of the software infrastructure provided by ISETBio project (isetbio.org), to which both DHB and NCP have contributed.

